# Interpretable Kolmogorov-Arnold Networks for Enzyme Commission Number Prediction

**DOI:** 10.1101/2025.01.30.633071

**Authors:** Louis Dumontet, So-Ra Han, Axel Prouvost, Jun Hyuck Lee, Tae-Jin Oh, Mingon Kang

## Abstract

Accurate prediction of enzyme commission (EC) numbers remains a significant challenge in bioinformatics, limiting our ability to fully understand enzyme functions and their roles in biological processes. This paper presents the integration and evaluation of the next paradigm of deep learning architecture, named Kolmogorov-Arnold network (KAN), in state-of-the-art models for predicting EC numbers. KAN modules are incorporated into current state-of-the-art models to assess their impact on predictive performance. Additionally, we introduce a novel interpretation method, specifically designed for KANs, to identify relevant input features for a given prediction, addressing a current limitation in KANs. Our evaluation demonstrates that the integration of KANs significantly enhances predictive performance compared to the state-of-the-art deep learning models, with up to a 4.35% increase in micro-averaged ***F***_**1**_ score and a 4.1% increase in macro-averaged ***F***_**1**_ score. Moreover, our novel interpretation method not only enhances the predictions’ trustworthiness but also facilitates the discovery of motif sites within enzyme sequences. This innovative approach provides deeper insights into enzyme functionality and highlights potential new targets for research. The results underscore KANs’ effectiveness in improving enzymatic classification and advancing our understanding of enzyme structures and functions. The open-source code is publicly available at: https://github.com/datax-lab/kan_ecnumber.

## 1 Introduction

Identification of protein chemical properties is essential for various biomedical applications, including protein-protein interaction prediction [1], diagnosing neurodegenerative diseases [2], and developing pharmaceuticals [3]. The protein chemical properties are categorized by Enzyme Commission (EC) numbers. This EC number system is critical for characterizing unknown enzymes which catalyze various commercial processes, such as pharmaceutical biosynthesis, food production, and bioremediation [4]. EC numbers categorize biological roles of enzymes in catalyzing major chemical reactions to biological processes across all organisms [5]. This classification system is organized into hierarchical levels: a class, a subclass, a sub-subclass, and a serial number (e.g., EC 1.2.3.4) [6–8].

Deep learning has significantly enhanced the automation of EC number predictions, enabling biologists to collaborate with advanced models to reduce the need for extensive and potentially unnecessary biological experiments. State-of-the-art deep learning models such as CLEAN [9], DeepECtransformer [10], DeepEC [11], ECPICK [12], ifDEEPre [13], HDMLF [14], and HECNet [15], have been developed to be increasingly effective when predicting enzymatic functions. This allows researchers to quickly annotate new sequences with high confidence, facilitating the functional characterization of enzymes.

Model interpretability is a crucial aspect of deep learning as it facilitates its understanding and assesses its robustness and trustworthiness. Interpretation of deep learning models for EC number predictions could provide insights into uncovering patterns in enzyme sequences which could deepen the understanding of the enzymatic activities and guide future research efforts. DeepECtransformer showed a preliminary approach to identify active site and binding residues in EC number prediction [10]. ECPICK introduced an interpretation method that enhances the trustworthiness of its predictions and uncovers potential new motif sites in enzyme sequences [12].

Recently, Kolmogorov-Arnold Networks (KANs) have been highlighted as a promising alternative to multilayer perceptrons (MLPs) [16]. In KANs, weight parameters are replaced by learnable functions, parameterized as splines, based on the eponymous theorem [17]. KANs combine the strengths of splines and MLPs’ compositional layer structures by optimizing both feature learning and univariate function approximation. The resulting architecture shows better performance than MLPs for simple tasks, such as predicting multi-variable functions or solving partial differential equations [16]. KANs have demonstrated their effectiveness and interpretability for low-dimensional problems and non-stochastic datasets. However, to fully leverage their potential for high-dimensional challenges, including protein sequence analysis, further exploration is needed. While KANs offer excellent interpretability for simpler tasks, developing advanced methods for interpreting these models in complex, high-dimensional contexts will be crucial for their successful application in real-world biological analyses.

In this study, our hypotheses are (Fig. 1): (1) KANs could significantly enhance performances when integrated in state-of-the-art models for EC number prediction and (2) KANs could increase model interpretability and further enhance biological understanding of the parts of enzyme sequences that are relevant to a predicted EC number. To prove these hypotheses, we explore the applicability of KANs, as an alternative to MLPs, for high-dimensional protein sequence analysis, and propose a novel interpretation approach for KANs. We also explore KAN architecture optimization strategies: pruning and grid extension. In this study, we specifically focus on EC number prediction with three digits of the EC number, which characterizes the physical properties of enzymes. The contributions of this study include (1) the first introduction of KANs to the real-world application of protein sequence analysis, (2) significant improvement of the predictive performance with KANs for EC number predictions, (3) the introduction of tuning strategies such as pruning and grid extension for high-dimensional data, and (4) the development of a novel interpretation strategy for KANs.

**Fig. 1.**
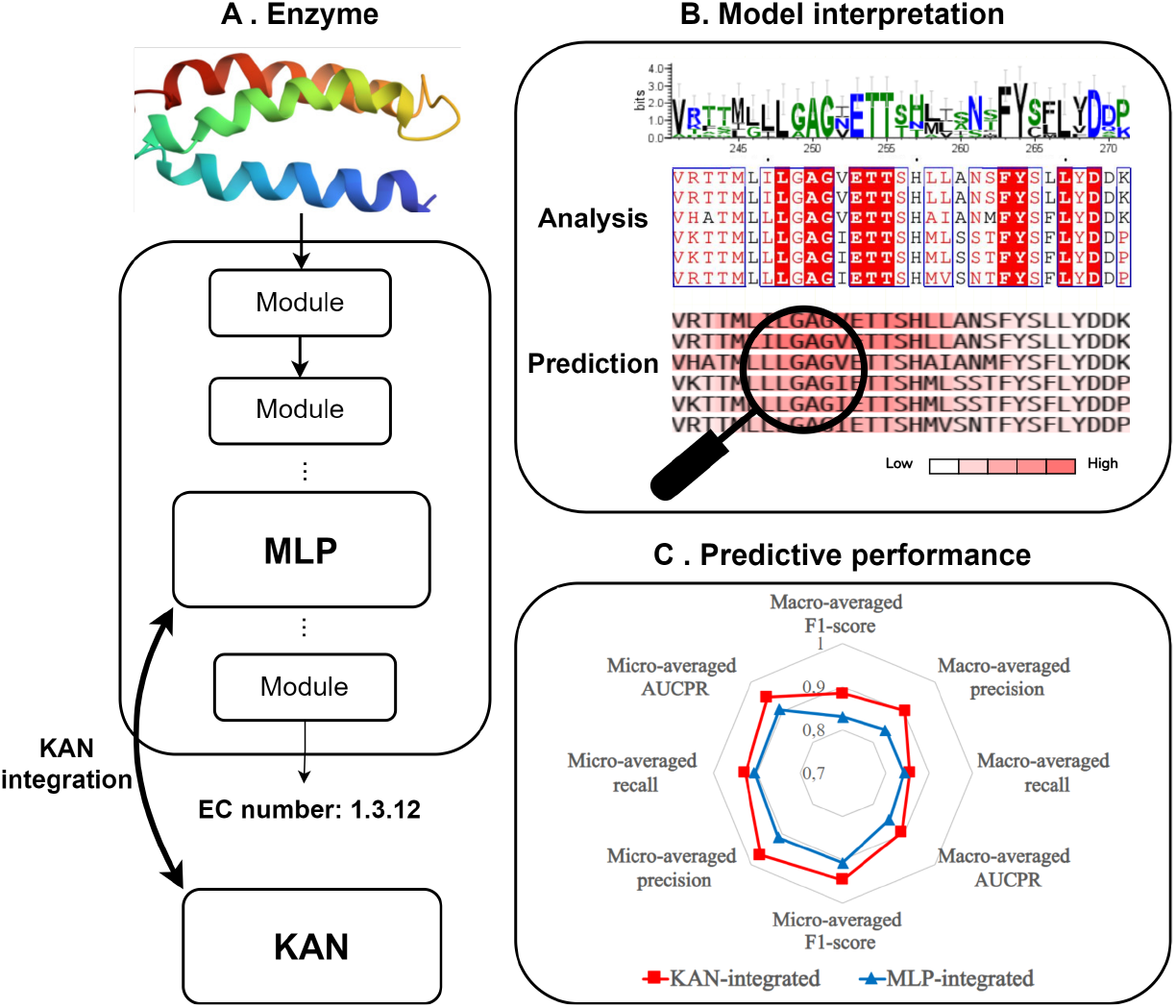
Overview of the study: (A) KANs are integrated in state-of-the-art EC number prediction models by replacing the MLPs. (B) The resulting models are interpretable as they can identify motif sites in enzyme amino-acid sequences, as in [18, 19]. (C) State-of-the-art models with KAN yield enhanced performances compared to original ones.

This paper first reviews current state-of-the-art models in EC number prediction. Then, we elucidate the principles of KANs in the methods section, including their architecture and the pruning and grid extension tuning strategies. We then introduce our novel interpretation method for KANs, demonstrating its utility in enhancing model transparency and performance. This is followed by a description of the integration of KAN modules into existing state-of-the-art models. In the experiment section, we provide an in-depth evaluation of our method, including dataset descriptions, performance comparisons between original and KAN-integrated models, analyses of the interpretation results, and the exploration of the tuning strategies. Finally, the conclusion summarizes the overall impact of this study, suggesting directions for future research and further application of KANs in bioinformatics.

## 2 Current deep learning models for EC number prediction

Deep learning models for EC number prediction can be broadly categorized into three main types: CNN-based, attention-based, and large language models (LLMs). CNN-based models leverage convolutional layers to extract hierarchical features from enzyme sequences or structures, effectively capturing local patterns and spatial information. Examples include DeepEC [11], ECPICK [12], and ECNet [20]. Attention-based models utilize the self-attention mechanism to improve the models’ ability to capture long-range dependencies and context. Notable examples are BEC-Pred [21] and DeepECtransformer [10]. LLMs, on the other hand, employ extensive pre-training on large datasets to understand protein sequences, which leverages contextual embeddings to predict protein functions and structures. Examples include CLEAN [9], AlphaFold [22], and ProtTrans [23]. Each category offers unique strengths and applications, reflecting the diverse approaches in state-of-the-art EC number prediction.

DeepEC is a representative example of CNN-based models. At its core, DeepEC employs convolutional layers to capture patterns within protein sequences, which are crucial for identifying enzymatic functions. Its architecture includes multiple convolutional layers that extract hierarchical features, followed by pooling layers to reduce dimensionality and emphasize the most relevant information. This is then complemented by MLP layers which integrate the extracted features and perform the final classification to predict the EC numbers. This approach enables DeepEC to leverage the strengths of convolutional neural networks in handling complex and high-dimensional enzyme sequences.

The regular transformer architecture [24] serves as a key example of attention-based models as several state-of-the-art methods heavily rely upon this architecture [10, 21]. It consists of multiple self-attention layers and MLPs. The regular transformer architecture leverages its ability to handle complex sequence data which provides robust predictions based on comprehensive contextual information.

CLEAN stands out as a prominent example of LLMs as it builds upon the ESM-1b model [25] as its foundation, which is an LLM specifically designed for protein sequences. To make its predictions, CLEAN integrates MLP layers on top of ESM-1b. This combination allows CLEAN to leverage ESM-1b’s deep contextual embeddings while utilizing the MLP layers to output a representation of the input sequence. CLEAN employs contrastive learning techniques, which train the model to encode protein sequences such that enzymes with similar activities are represented closely in the embedding space, while those with different activities are positioned farther apart.

## 3 Methods

This section provides details about the fundamentals of the KAN architecture and its architecture-tuning approaches: pruning and grid extension. Then, we introduce our novel interpretation strategy which identifies relevant input features to the predictions. Finally, we explain how KANs can be integrated into state-of-the-art models.

### 3.1 Architecture of Kolmogorov-Arnold Networks

KANs, like MLPs, are fully connected feed-forward networks which leverage dense connectivity to capture complex and non-linear relationships for inferring desired outcomes. MLPs learn weight parameters and use fixed activation functions, whereas KANs do not learn weights; rather, they replace fixed activation functions with learnable activation functions (Fig. 2). According to the Kolmogorov-Arnold representation theorem, any complex high-dimensional function can be represented by a polynomial number of univariate functions [17]. Thus, KAN, which is a neural network with compositions of univariate functions, can effectively model complex and high-dimensional functions. In KANs, each layer of a neural network represents a component of the compositions, which collectively model relevant relationships within the data.

**Fig. 2.**
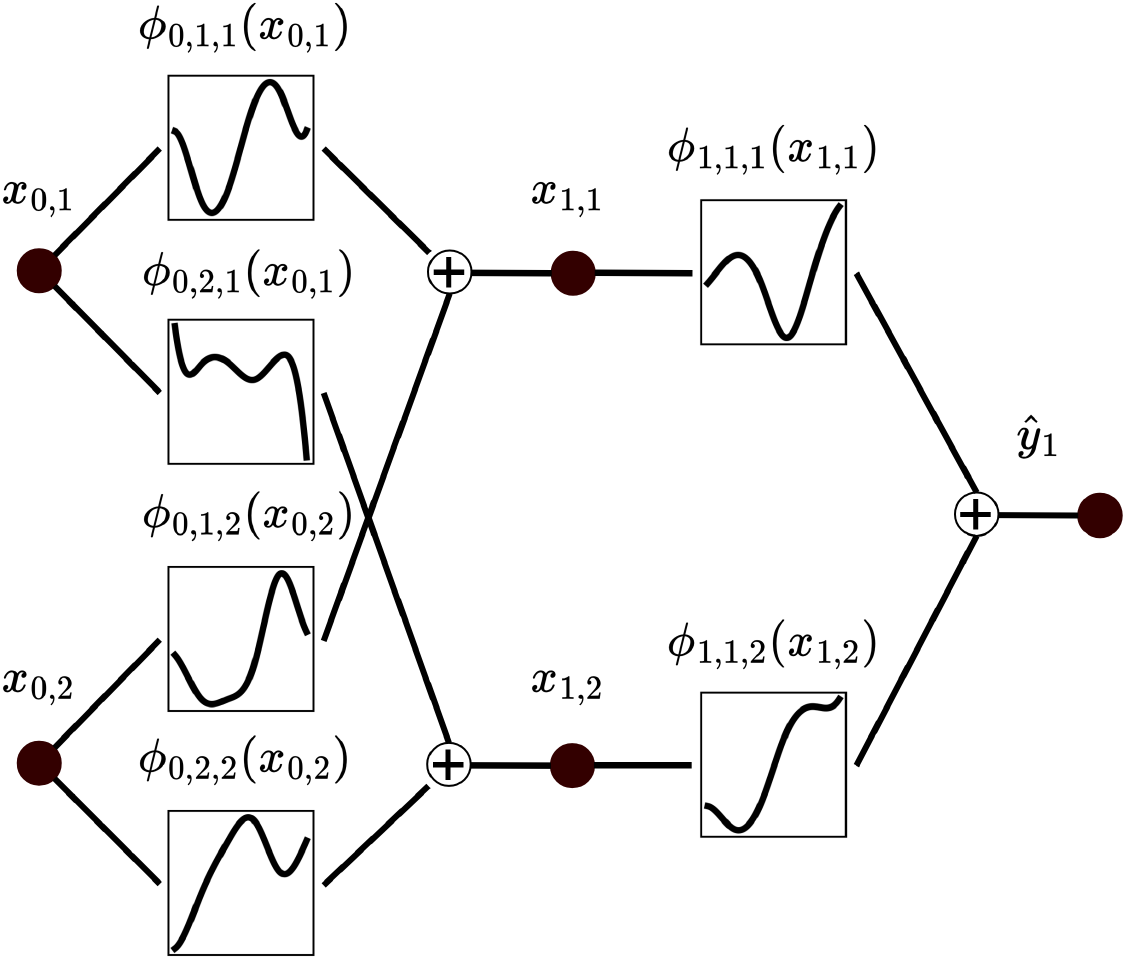
Representation of a two-layer KAN of dimensions 2 and 1, with an input of dimension 2: (*x*_0,1_, *x*_0,2_). *ϕ*_*l*,*j*,*i*_ is the activation function between the *i*^*th*^ node of the *l*^*th*^ layer, and the *j*^*th*^ node of the *l* + 1^*th*^ layer. For any layer, each node is connected to all the nodes of the next layer.

A KAN (𝒦 : **x** → **ŷ**) is formulated as:

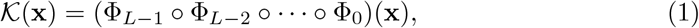

where *L* is the number of layers, Φ_*l*_ is a function matrix, and ° is the composition operator, such as (Φ_1_ ° Φ_2_)(*x*) = Φ_1_(Φ_2_(*x*)). The output **ŷ** is obtained by computing the composition of *L* functions (Φ_*l*_) from the input **x**. The function matrix, Φ_*l*_, is a set of activation functions *ϕ*_*l*,*j*,*i*_, connecting the *i*^*th*^ neuron in the *l*^*th*^ layer to the *j*^*th*^ neuron in the *l* + 1^*th*^ layer, such as:

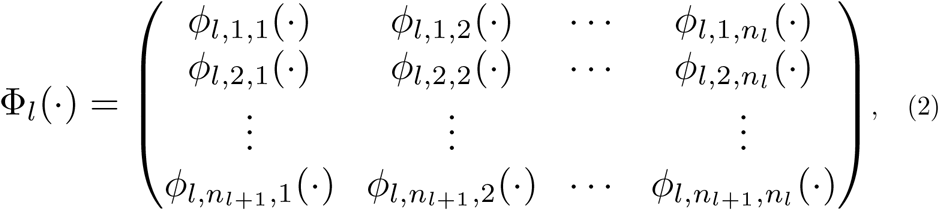

where the layer sizes are [*n*_0_, *n*_1_, …, *n*_*L−*1_].

For the activation function, *ϕ*_*l*,*j*,*i*_, the input value (*x*_*l*,*i*_) is called pre-activation, and the output of the activation function 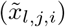 is called post-activation, such as:

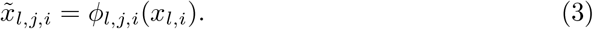

Then, the pre-activation value for the *j*^*th*^ neuron in the next layer is *x*_*l*+1,*j*_, which is the sum of post-activations from the previous layer 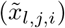, such as:

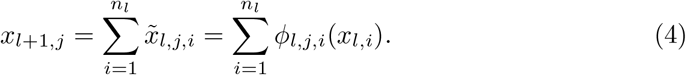

A post-activation (*ϕ*_*l*,*j*,*i*_(*x*_*l*,*i*_)) is computed as:

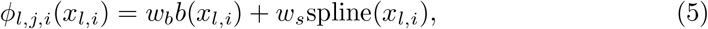

where *w*_*b*_ and *w*_*s*_ are trainable parameters that control overall magnitude of the activation function, and the function *b* is the SiLU function (i.e., 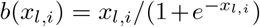) which allows the networks to bypass one or more layers, similarly to residual connections in MLPs. The spline function is defined as:

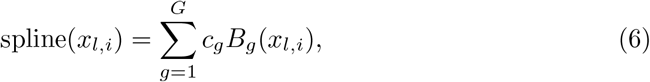

where *B*_*g*_s are piecewise polynomial functions called B-splines, and *c*_*g*_s are trainable parameters. The resulting linear combination is a spline of order *k*, which is defined on a specific interval of *G* grid-points.

The complexity of a KAN with *L* layers of width *N* is *O*((*G*+*k*)*N* ^2^*L*), which would be *O*(*N* ^2^*L*) for a same dimension MLP. However, KAN has been proven to require much smaller networks [16].

### 3.2 Architecture-tuning strategies with KAN

In this section, we present two architecture-tuning strategies for KANs: pruning and grid extension. Pruning is a sparsification technique to enhance the efficiency of KANs by systematically reducing the network’s parameters. On the other hand, grid extension is a strategy to accelerate the training process of KANs, thus optimizing computational resources. Each strategy is explored with detailed explanations, highlighting their respective contributions to improving the performance and efficiency of KANs.

Pruning reduces the number of parameters of the network by suppressing irrelevant connections (Fig. 3A). The goal of pruning is to make the network sparser, which reduces computational costs and memory usage. Neurons are considered irrelevant if the maximum value between their incoming and outgoing scores is not higher than a threshold (*θ*). Mathematically, the *i*^*th*^ neuron of the *l*^*th*^ layer is pruned if:

**Fig. 3.**
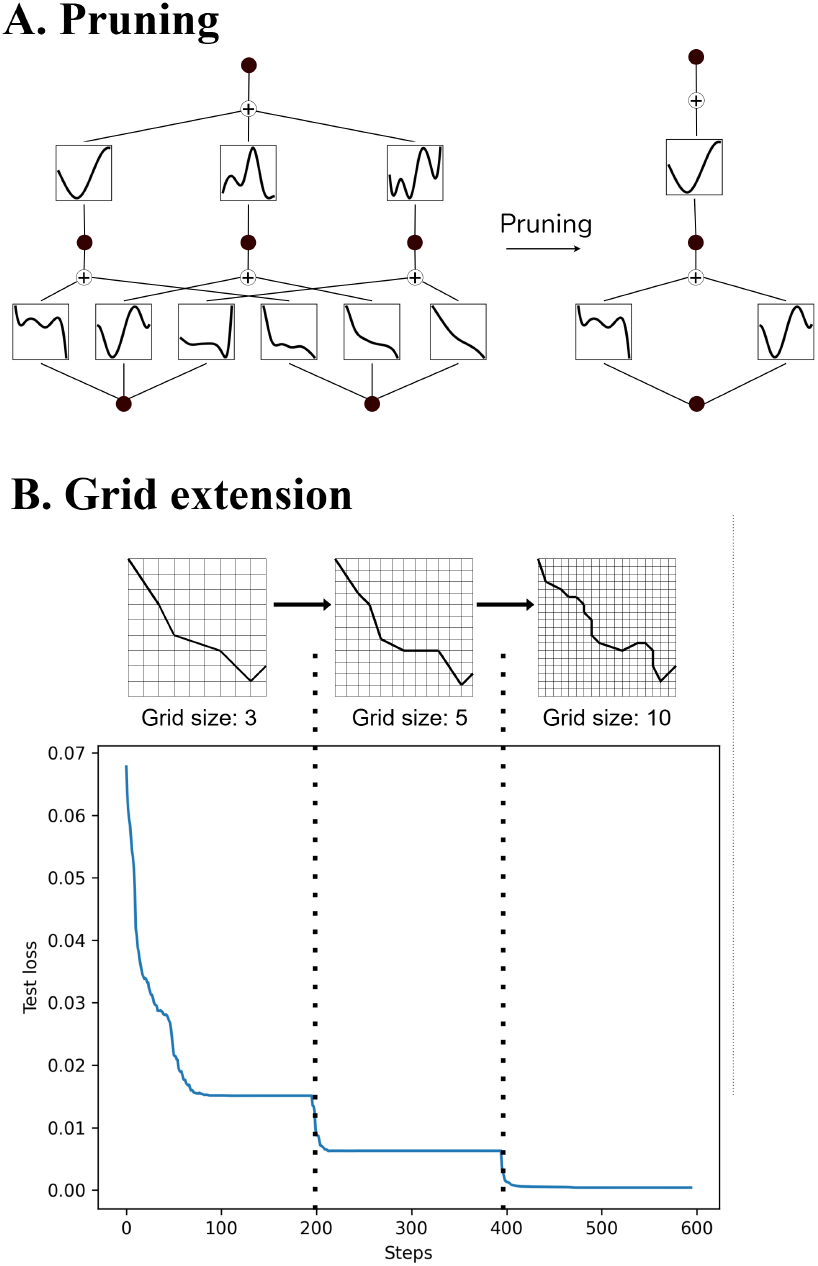
Architecture-tuning strategies for KAN. (A) A KAN before and after pruning. Pruning removes the irrelevant connections of the network, allowing the model to contain less parameters and to perform faster predictions without retraining it. (B) Grid extension changes the grid resolution during training, which makes the training process more efficient.

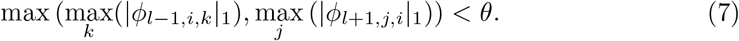

Pruning does not require the model to be retrained, which makes it a cost-effective sparsification technique that manages the trade-off between efficiency and predictive performance.

Grid extension is a tuning strategy that adjusts the grid resolutions for activation functions during the training process. To enhance training efficiency, a coarse grid can be extended to a finer grid during the training process, with the coarse grid enabling more efficient training and the fine grid providing more accurate function representations.(Fig. 3B) Consequently, grid extension combines the efficiency of coarse grids with the accuracy of fine grids [16].

Let *G*_1_ be the number of points of the initial grid and 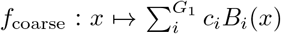 a spline. To extend the initial grid to a finer grid with *G*_2_ points and a spline 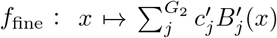, the 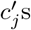 are obtained by minimizing the distance between *f*_coarse_ and *f*_fine_ using the least square algorithm.

### 3.3 KAN Interpretability

We propose a new interpretation strategy that identifies a feature set which is relevant to the prediction and, address a critical gap, as such interpretation methods do not exist for KANs. This strategy identifies relevant amino acids in the protein sequence that contribute to a given prediction, thereby offering insights into biological processes of enzyme functions.

In a KAN with L layers of dimensions [*n*_0_, *n*_1_, …, *n*_*L−*1_], a posterior probability (*ŷ*_*c*_) is computed by applying the sigmoid function on the last activation value (*s*_*c*_) for the class c, such as:

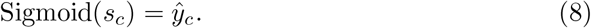

The relative contribution to the prediction of each neuron in the *L −* 1^*th*^ layer is represented by a vector, **s**_**L***−***1**_ = {*s*_*L−*1,*j*_; 1 ≤ *j* ≤ *n*_*L−*1_}, as:

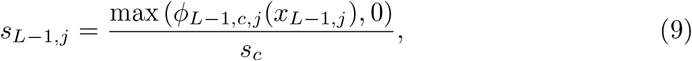

where *ϕ*_*L−*1,*j*,*i*_ is the activation function between the *i*^*th*^ node of the *L* − 1^*th*^ layer and the *j*^*th*^ node of the *L*^*th*^ layer.

For any layer *l* between 1 and *L −* 2, *x*_*l*,*i*_ is the pre-activation value of *ϕ*_*l*,*j*,*i*_ as defined in (4). We define **s**_**l**_ = {*s*_*l*,*j*_; 1 ≤ *j* ≤ *n*_*l*_} as the contribution scores for each neuron of the *l*^*th*^ layer, which is computed as:

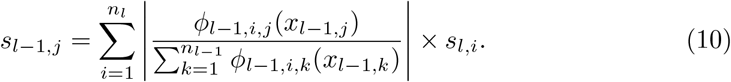

This operation (Eq. 10) is repeated on each layer, starting with the *L −* 2^*th*^ layer to the first layer, thus obtaining the contribution scores **s**_**1**_ of the input **x**.

### 3.4 Integration of KAN in state-of-the-art models

In this study, we integrated KAN layers into state-of-the-art models in three categories: CNN-based models, attention-based models, and LLMs for protein sequences, to perform a thorough assessment of their impact and effectiveness covering the most advanced and high-performing architectures available. For the CNN-based model, we integrated KAN layers in the DeepEC architecture (Fig. 4A). We replaced the three MLP layers by two KAN layers of 512 and 229 nodes, respectively, with a grid size of three and a spline order of three, obtained through fine-tuning the hyper-parameters of KAN-integrated DeepEC.

**Fig. 4.**
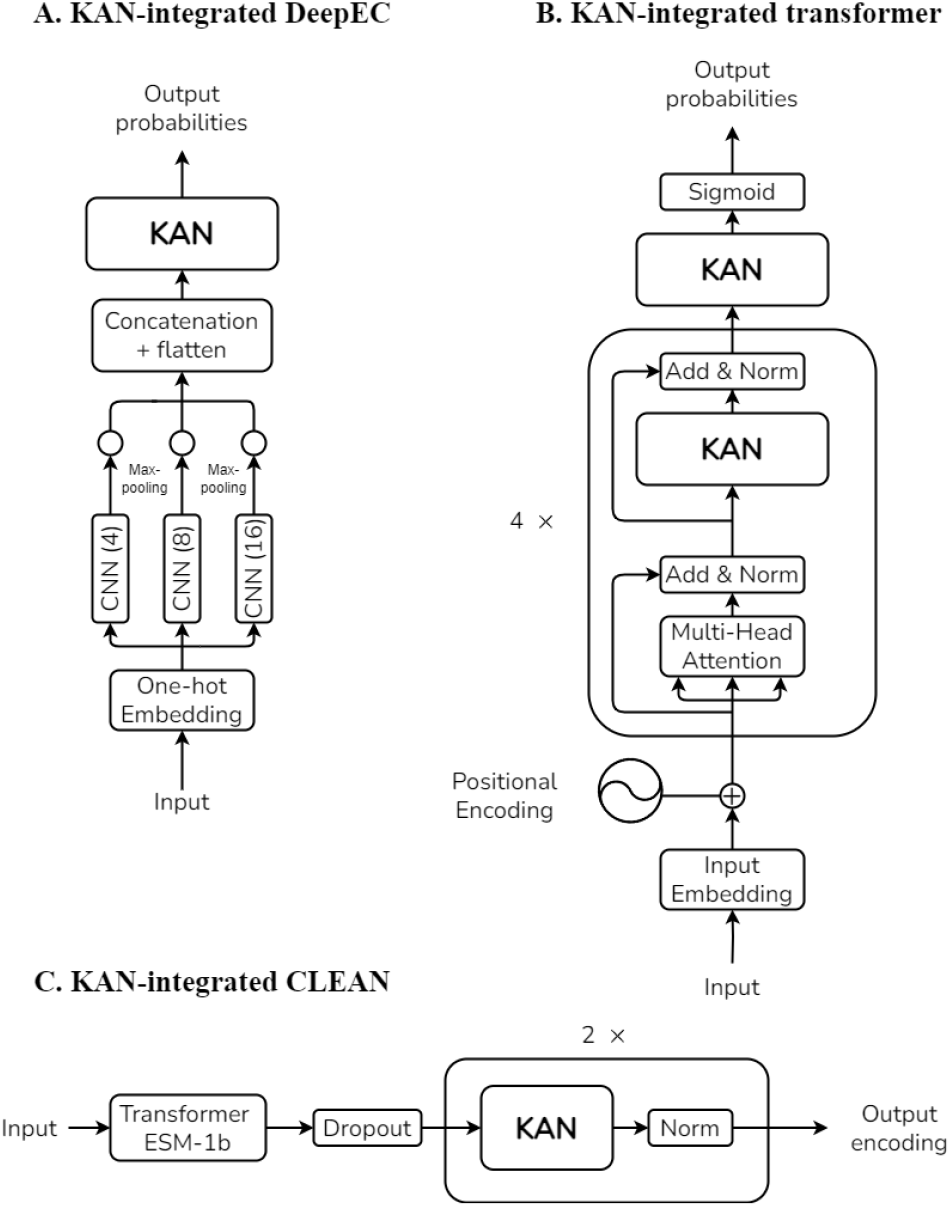
Integration of KAN into state-of-the-art models improves their predictive performances. In DeepEC (A) KAN layers replace MLP layers after the three parallel CNNs. In the regular transformer (B), both the neural networks following the multi-headed attention module, and the output layer are replaced by KAN layers. In CLEAN (C), KAN layers replace the MLP layers after the ESM-1b transformer.

For the attention-based model, regular transformers were combined with KANs by replacing the MLP layers after the multi-headed attention modules and replacing the classification MLP layers (Fig. 4B) [24]. We considered two commonly-used dimensions for transformers, 128 and 256. After the hyper-parameter tuning for the 128-dimensional transformer, we replaced the three MLP layers following the multiattention head module by a KAN layer comprising 128 nodes, with a grid size of five and a spline order of one. For the 256-dimensional transformer, we replaced the same three MLP layers by a KAN layer containing 256 nodes, with a grid size of three and a spline order of three. On the other hand, the replaced classification layers have a fixed size of 229 nodes as we classify enzymes amongst 229 EC numbers.

For the LLM model, we enhanced the CLEAN architecture by replacing its three MLP layers with two KAN layers, while maintaining dropout and layer normalization as proposed in the original model. Following the hyper-parameter optimization, the KAN-integrated CLEAN model was configured with two KAN layers of 512 and 229 nodes, a grid size of three, a spline order of three, and a dropout rate of 0.3 (Fig. 4C).

## 4 Experimental Results

### 4.1 Dataset

For this study, we utilized the protein sequence data collected from Swiss-Prot [26] and Protein Data Bank (PDB) [19], published before September 2022. The enzymes are distributed across 229 EC numbers which are each associated with at least 10 samples in the dataset. We preserved protein sequences, containing up to 1,000 amino acids and non-redundant sequences. Each sequence is associated with at least one EC number. The resulting dataset contains more than 200,000 protein sequences of which approximately two-thirds come from Swiss-Prot and one-third comes from PDB.

### 4.2 The integration of KANs enhances predictive performance

We compared the predictive performance between the KAN-integrated models and their original counterparts. We split the dataset into training (80%), validation (10%), and test (10%) sets using stratified random sampling to preserve class ratios. Specifically, the sample sizes were approximately 160,000 for the training set, 20,000 for the validation set, and 20,000 for the test set. We optimized the models with the training dataset, while the hyper-parameters were fine-tuned with the Optuna framework [27] using the validation dataset. Then, we assessed the performance of the models on the test dataset. We repeated the experiment ten times for reproducibility.

We computed the micro and macro-averaged precision, recall, and *F*_1_ score to evaluate the models. For each model, we selected the threshold for the final discriminative function that maximizes the macro-averaged *F*_1_ score on the validation set. Then, we computed the evaluation metrics on the test set using these thresholds.

Across all the state-of-the-art methods, KAN-integrated models demonstrated significant improvements over the original models, achieving up to a 4.35% increase in micro-averaged *F*_1_ score and up to a 4.1% increase in macro-averaged *F*_1_ score (Table 1). Specifically, KAN-integrated DeepEC achieved a macro-averaged *F*_1_ score of 86.4% *±* 2.1%, outperforming the original DeepEC model’s macro-averaged *F*_1_ score of 83% *±* 2.4%. The KAN-integrated transformers showed superiority over regular transformers in both dimensions 128 and 256, with macro-averaged *F*_1_ scores of 79.5% *±* 2% and 86.8%*±*1.9% respectively, against 78.4% *±* 2.1% and 85.6% *±* 2% for the original transformers. The KAN-integrated CLEAN also achieved a higher macro-averaged *F*_1_ score of 90.5% ± 0.8%, compared to 88.7% ± 0.8% for the original model. The statistically significant improvements with the KAN-integrated models were confirmed by Wilcoxon signed-rank tests across all the models (*p <* .05).

**Table 1.** Performance comparison between state-of-the-art model and their KAN-integrated counterparts. A single/double asterisk indicates *p <* .05 (*) and *<* .01 (**) for the Wilcoxon signed-rank test, respectively.

| 3-8 |  | Micro-averaged |  |  | Macro-averaged |  |  |
| --- | --- | --- | --- | --- | --- | --- | --- |
| | | $F_1$ Score (%) | Precision (%) | Recall (%) | $F_1$ Score (%) | Precision (%) | Recall (%) |
| DeepEC | MLP | 90.80 ± 1.04 | 91.13 ± 1.13 | 90.47 ± 1.20 | 83.02 ± 2.44 | 84.00 ± 2.30 | 84.24 ± 2.29 |
|  | KAN | **94.75 ± 0.90\*\*** | **96.97 ± 0.90** | **92.64 ± 1.42** | **86.43 ± 2.10\*\*** | **90.37 ± 2.48** | **84.51 ± 2.00** |
| Transformer 128 | MLP | 88.59 ± 1.50 | **92.47 ± 0.97** | 85.02 ± 2.04 | 78.44 ± 2.15 | **87.95 ± 1.73** | 73.35 ± 2.40 |
| | KAN | **89.39 ± 1.16\*** | 91.58 ± 0.99 | **87.31 ± 1.71** | **79.51 ± 2.00\*** | 81.42 ± 1.78 | **77.09 ± 2.18** |
| Transformer 256 | MLP | 91.98 ± 1.73 | **94.83 ± 1.28** | 89.30 ± 2.17 | 85.59 ± 2.04 | **91.96 ± 1.32** | 81.86 ± 2.42 |
| | KAN | **92.29 ± 1.33\*** | 93.73 ± 1.11 | **90.91 ± 1.49** | **86.76 ± 1.93\*** | 91.01 ± 2.24 | **84.36 ± 1.79** |
| CLEAN | MLP | 92.46 ± 1.78 | 92.80 ± 1.65 | 92.13 ± 1.99 | 88.66 ± 0.79 | 87.81 ± 0.84 | 93.19 ± 1.16 |
|  | KAN | **93.73 ± 1.52\*\*** | **94.48 ± 1.38** | **93.00 ± 1.71** | **90.46 ± 0.84\*\*** | **90.37 ± 0.81** | **93.52 ± 1.21** |

The improvements in predictive performance are positively correlated to the proportion of MLPs in the original models. The original architecture of DeepEC comprises MLPs that take 97.5% of learnable parameters. However, MLPs account for 88.5%, 69.4%, and 71% of the learnable parameters in CLEAN, and transformers with dimensions 128 and 256, respectively. We also noted that KAN-integrated models produce more consistent performance across the experiments. The standard deviations decreased by 19.2% for the micro-averaged *F*_1_ score and by 7.6% for the macro-averaged *F*_1_ score across the experiments (Table 1). Moreover, KAN-integrated transformers produced more balanced performance between precision and recall than the original transformers.

### 4.3 KANs identify existing motif sites by the proposed interpretation strategy

We verified the proposed KAN interpretation strategy that identifies relevant amino acids, comparing them with well-reported motif sites on enzyme sequences. We computed a contribution score for each amino acid to represent its impact in the model’s predictions from a given protein sequence. We focused on the KAN-integrated DeepEC model as it showed the most significant performance improvement.

The proposed interpretation strategy computed the intermediate scores of the activation maps, which are the input of the KAN module. To map the intermediate scores (**s** = {*s*_*q*_; 1 ≤ *q* ≤ 384}) to the contribution scores of the protein sequence (*γ* = {*γ*_*q*_; 1 ≤ *q* ≤ 1000}), we identified the segments of the sequence that produced the 384-dimensional input vector to the KAN module (**v** = {*v*_*j*_; 1 ≤ *j* ≤ 384}). Each input value *v*_*j*_ is the maximum value of the corresponding activation map **f**_**j**_, and *℩*_*j*_ is its index, such as:

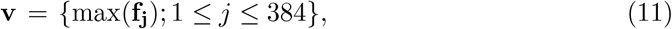

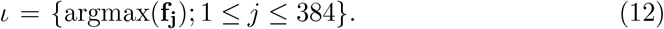

We grouped the intermediate scores from the activation maps of size *z* (i.e., kernel size: 4, 8, or 16) and index *i* in a set 𝒥_*i*,*z*_. Mathematically, 𝒥_*i*,*z*_ is:

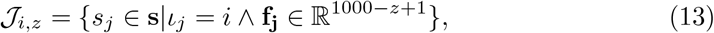

where ∧ is the logical AND operator. Thus, the intermediate scores 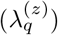 of the *q*^*th*^ amino acid from the activation maps of size *z* is:

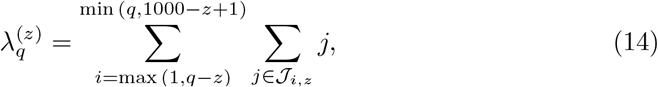

where the range from max (1, *q* − *z*) to min (*q*, 1000 − *z* + 1) represents all the possible indices of an activation map of size *z*, which is computed from the *q*^*th*^ amino acid. To compute the contribution score (*γ*_*q*_) of the *q*^*th*^ amino acid in the input protein sequence, we summed the intermediates scores of activation maps of size 4, 8, and 16 computed from the *q*^*th*^ amino acid, such as:

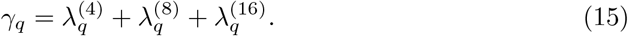

For the validation of the proposed interpretation strategy, we considered enzyme sequences from Cytochrome P450 (CYP) (CYP106A2 family [EC 1.14.15] and CYP7B1 family [EC 1.14.14]), whose biological functions are well-reported in various organisms, from bacteria to mammals [28]. CYPs belong to a super-family of enzymes that have been extensively studied, and widely utilized in the pharmaceutical industry and in clinical and disease-related medicine [29]. CYPs are monooxygenases that catalyze the incorporation of a single oxygen atom into substrates. For example, CYP106A2 and CYP7B1 enzymes perform similar biological functions in different organisms (e.g., bacteria and human). The bacterial CYP106A2 group plays a crucial role in attaching a hydroxyl group to steroid structures. Whereas, the human CYP7B1 group is involved in the metabolism of endogenous oxysterols and steroid hormones, including neurosteroids, in eukaryotic cells. Despite these functional differences, both CYP106A2 and CYP7B1 share the same first and second digits in their EC numbers, indicating that they belong to the same general category of oxidoreductases (EC 1) that act on paired donors and involve the incorporation or reduction of molecular oxygen (EC 1.14).

We performed the model interpretation on 13 sequences from the bacterial CYP106A2 enzyme family. Specifically, we considered 5XNT [30] and 4YT3 [31] from the Protein Data Bank (PDB), along with 11 protein sequences from Swiss-Prot that share over 90% sequence similarity with 5XNT and 4YT3. We computed the contribution scores of amino acids for each protein sequence using the KAN interpretation method and ECPICK, which is the current state-of-the-art interpretation strategy to identify motif sites in enzyme sequences [12]. Both KAN and ECPICK correctly predicted all of the protein sequences as belonging to the 1.14.15 class.

The 13 protein sequences were aligned by a multiple sequence alignment (MSA) tool (e.g., Clustal Omega [32]) for graphical comparison. Conserved amino acids were identified by ESPript3 [33], colored in red. Fig. 5A illustrates the corresponding contribution scores of KAN and ECPICK along with motif sites (e.g., oxygen-binding, EXXR, and heme-binding domains) in black boxes and substrate recognition sites (SRS 1-6) in green boxes, which are key regions relevant to the enzyme function. High contribution scores are indicated in dark red, with lower scores gradually transitioning to white along the gradient scale.

**Fig. 5.**
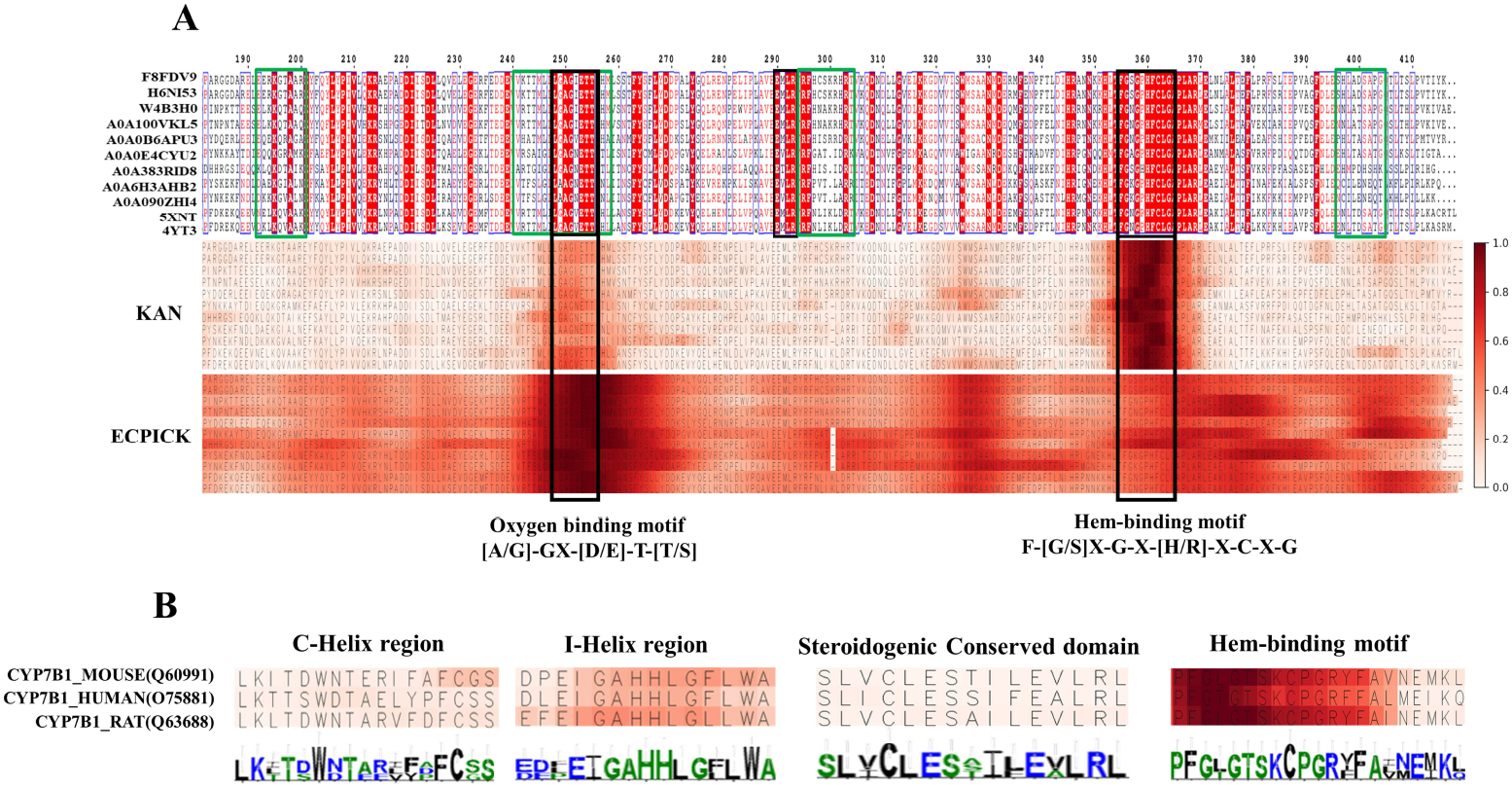
The model identifies significant amino acids contributing to the prediction, demonstrating trustworthiness in its predictions and discovering motif sites. High contribution scores are indicated in dark red, with lower scores gradually transitioning to white along the gradient scale. (A) A partial sequence, spanning positions 182 to 418, from the CYP106A2 family is displayed. This segment was selected for detailed analysis due to its significant role in CYP106A2. Motif sites are outlined in black boxes and substrate recognition sites (SRS 1-6) in green boxes. KAN interpretation highlights the oxygen-binding site and the hem-binding motif, which are primary active sites within the CYP106A2 group. On the other hand, ECPICK only recognizes the oxygen-binding motif site. (B) Visualizes the four motif sites in the protein sequences of the CYP7B1 family with contribution scores computed by the proposed KAN interpretation, which emphasizes the central role of these regions in predicting the enzymatic function (EC 1.14.14).

The proposed KAN interpretation method successfully identified both the oxygen binding motif site and hem-binding motif sites of the bacterial CYP106A2 family. The identification of the binding motif sites aligns with established biological knowledge, which determines the primary and secondary levels of the given EC number (e.g., 1.14). However, ECPICK did not recognize the hem-binding motif site, which is crucial for determining the CYP’s enzyme function. The EXXR motif site was not identified by either model, as EXXR may not be sufficiently discriminative for the given EC number (e.g., 1.14).

We also analyzed the proposed strategy within the CYP7B1 group across the three organisms: mice, rats, and humans. KAN-integrated DeepEC accurately predicted the enzyme functions of the protein sequences (i.e., 1.14.14). Moreover, the contribution scores effectively highlighted the essential motif sites of this CYP enzyme family (Fig. 5C). Specifically, the hem-binding site was highly scored across the three species. The I-helix, which contains the oxygen-binding site, and the C-helix regions were identified by the proposed interpretation, whereas the steroidogenic conserved domain was assigned low contribution scores.

High contribution scores were observed in the essential motif sites on both the CYP106A2 and CYP7B1 families. The identified sequential patterns aligned well with conserved domains or existing motif sites. This analysis was achieved without the time-consuming computational processes typically required for sequence similarity and secondary structure comparison. The proposed interpretation strategy provides trustworthiness in prediction by identifying existing motif sites related to the enzyme function and could potentially discover unknown motif sites within enzyme sequences.

### 4.4 Pruning and grid extension for architecture optimization

We conducted an additional experiment to evaluate the pruning and grid-extension strategies for KANs. We tested pruning to remove irrelevant connections of the KAN-integrated DeepEC model. We initiated this experiment by training a four-layer KAN of dimensions 512, 1024, 512, and 229 in KAN-integrated DeepEC. After training the model, we applied pruning as defined in (7) with varying thresholds (*θ*). Fig. 6 illustrates the macro-averaged *F*_1_ scores of the pruned models in relation to the number of parameters controlled by varying thresholds.

**Fig. 6.**
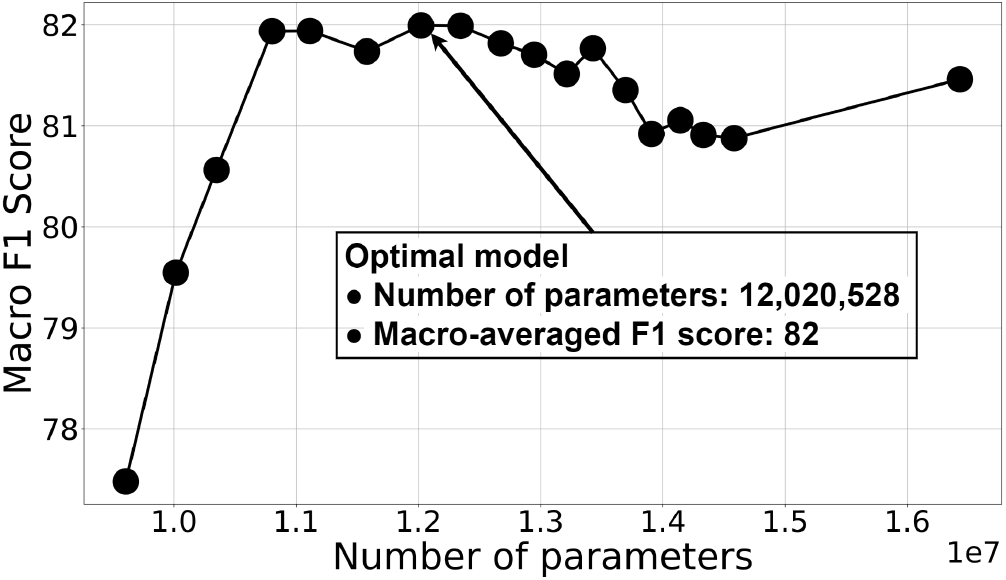
Pruning of a KAN-integrated DeepEC. A KAN model is pruned with varying thresholds. The number of parameters thus decreases incrementally as the thresholds increase. We observed that the best performance is yielded by a pruned model. The unpruned model contains 36.68% more parameters than the best-performing model.

The best model achieved a macro-averaged *F*_1_ score of 82% and contains 12,020,528 parameters, whereas the unpruned model yielded a macro-averaged *F*_1_ score of 81.46% with 16,429,184 parameters. The unpruned model comprises 36.68% more parameters than the pruned model. Hence, the removal of the connections increases the efficiency and robustness of the model as well as its predictive performance. Pruning reduces the size of the network without retraining the model. However, the improvement in predictive performance and efficiency highly depends on the unpruned model’s architecture.

In the experiment, the tuning with grid extension did not improve the predictive performance. This experiment result implies potential limitations of grid extension in handling high-dimensional and noisy real-world data. Protein sequences, characterized by their inherent complexity and variability, may present challenges that were not accounted for in prior evaluations of grid extension, where only low-dimensional, non-stochastic datasets were considered [16]. The high dimensionality and noise in protein sequence data could have hindered the strategy’s effectiveness, suggesting a need for further investigation of new grid extension strategies.

## 5 Conclusion

In this study, we have investigated KANs’ potential for EC number prediction using protein sequence data. We evaluated the integration of KAN modules in state-of-the-art deep learning-based models for EC number prediction. We have also proposed a novel interpretation method for KANs that identifies relevant input features to the model’s predictions. The KANs’ significant improvement was demonstrated through various experiments, which highlight the potential of KANs in advancing towards more accurate enzyme function prediction models. Additionally, we evaluated our novel KAN interpretation method and ascertained its ability to identify motif sites in enzyme sequences, providing trustworthiness in the predictions and biological insights. The proposed interpretation method could guide future research efforts to uncover new potential motif sites. Finally, we have evaluated the tuning strategies, pruning and grid extension, for protein sequences. We have observed that pruning is applicable to this task and optimizes the KANs’ architecture without having to retrain the model, whereas the grid extension strategy is very sensitive to noise and unlikely to be effective for real-world data.

This study opens several paths for future research into KANs. Their ability to capture complex relationships within protein sequences could be further explored by integrating them with other protein sequence analysis methods. Moreover, the interpretability of KANs could be enhanced by coupling the proposed method with other explainability techniques, allowing for a better understanding of how specific motifs contribute to the prediction of enzyme functions. Overall, this study demonstrates the promising future of KANs and emphasizes the importance of model interpretability regarding protein sequence analysis. Future work in this area could significantly advance the development of robust and efficient models for handling complex biological tasks, which will contribute to the advancement of biological understanding.

## 6 Code availability

The open-source code is publicly available at: https://github.com/datax-lab/kanecnumber.

## 7 Acknowledgments

We acknowledge the supports from National Science Foundation Major Research Instrumentation (NSF MRI) (Grant#:2117941), Ministry of Oceans and Fisheries in the Republic of Korea (20200610, KOPRI Grant), and National Research Foundation of Korea (NRF) by the Korea government (MSIT) (RS-2024-00354012).

